# O-antigen diversification masks identification of highly pathogenic STEC O104:H4-like strains

**DOI:** 10.1101/2022.09.15.508078

**Authors:** Christina Lang, Angelika Fruth, Ian W. Campbell, Claire Jenkins, Peyton Smith, Nancy Strockbine, François-Xavier Weill, Ulrich Nübel, Yonatan H. Grad, Matthew K. Waldor, Antje Flieger

## Abstract

Shiga toxin-producing *E. coli* (STEC) can give rise to a range of clinical outcomes from diarrhea to the life-threatening systemic condition, hemolytic uremic syndrome (HUS). Although STEC O157:H7 is the serotype most frequently associated with HUS, a major outbreak of HUS occurred in 2011 in Germany, and was caused by a rare serotype, STEC O104:H4. Prior to 2011 and since the outbreak, STEC O104:H4 strains have only rarely been associated with human infections. From 2012 to 2020 intensified STEC surveillance was performed in Germany where subtyping of ∼8,000 clinical isolates by molecular methods including whole genome sequencing was carried out. A rare STEC serotype O181:H4 associated with HUS was identified and, like the STEC O104:H4 outbreak strain, this strain belongs to sequence type (ST) 678. Genomic and virulence comparisons revealed that the two strains are phylogenetically related and differ principally in the gene cluster encoding their respective lipopolysaccharide O-antigens but exhibit similar virulence phenotypes. In addition, five other serotypes belonging to ST678 from human clinical infection, such as OX13:H4, O127:H4, OgN-RKI9:H4, O131:H4, and O69:H4, were identified from diverse locations worldwide.

**Importance:** Our data suggest the high virulence ensemble of the STEC O104:H4 outbreak strain remains a global threat because genomically similar strains cause disease worldwide, but that horizontal acquisition of O-antigen gene clusters has diversified the O-antigens of strains belonging to ST678. Thus, identification of these highly pathogenic strains is masked by diverse and rare O-antigens, thereby confounding the interpretation of their potential risk.

## Introduction

Shiga toxin-producing *E. coli* (STEC) are food-borne pathogens responsible for a range of clinical syndromes from diarrhea to the life-threatening systemic condition, hemolytic uremic syndrome (HUS), a triad of thrombotic microangiopathy, thrombocytopenia, and acute renal injury (1). Classical STEC include the pathovar enterohemorrhagic *E. coli* (EHEC), such as strains of serotype O157:H7, which in addition to the Shiga toxin gene, contain the locus of enterocyte effacement (LEE) pathogenicity island coding for a virulence-associated type III secretion system and effectors (1). Historically, the classification of STEC strains into different serotypes has proven invaluable for epidemiology and risk profiling (2). In *E. coli*, the serotype is determined by a combination of O- and H-antigen types (see below) whereas the O group solely denotes the O-antigen. Globally, O157:H7 strains are most frequently associated with HUS, but furthermore strains belonging to O groups O26, O103, O111, and O145, have also been regularly linked to HUS development (2, 3). In addition, a very rare serotype gave rise to a major HUS outbreak in early summer 2011, when an O104:H4 strain caused more than 3,000 cases of diarrhea and 800 cases of HUS including 54 fatalities, predominantly in Germany (4–6).

The O104:H4 outbreak strain encodes an exceptional set of virulence features (5–9). Like other HUS-associated strains, the strain produces Shiga toxin (Stx), specifically the Stx2a variant. But unlike most *E. coli* strains causing HUS, this strain belongs to enteroaggregative *E. coli* (EAEC), acquired a *stx2a*-encoding phage, and lacks LEE. EAEC strains characteristically harbor a plasmid (pAA) encoding aggregative adhesion fimbriae (AAF); in this case AAF of type I (AAF/I) (5, 7). AAF are responsible for bacterial autoaggregation, the stacked-brick adhesion phenotype to host cells, and contribute to the inflammatory response (10, 11). Furthermore, the outbreak strain encodes the virulence-linked serine-protease autotransporters (SPATE), SepA, SigA, and Pic and harbors an additional plasmid encoding an extended spectrum beta-lactamase (ESBL) of the type CTX-M-15 (5, 11, 12). The O104:H4 outbreak strain along with other O104:H4 strains all belong to multi-locus sequence typing (MLST) sequence type (ST) ST678 and some possess *stx*, forming a distinct clade among EAEC (5, 7, 11). Despite the extensive outbreak in May/June of 2011 and associated wide distribution of the strain in affected regions, intensified molecular surveillance only uncovered relatively few O104:H4 cases in Germany after the outbreak dissipated by July 2011.

In *E. coli*, serotypes are determined by the composition of the lipopolysaccharide (LPS) O-antigen and the flagellar H-antigen, both important surface features of microorganisms that shape pathogen host interactions (13, 14). LPS forms a major structural component of the Gram-negative cell’s outer membrane and its most distal part is the O-antigen. The O-antigen is subject to strong selection pressure and is one of the most variable components of the bacterial cell (13). Typically, in *E. coli* the O-antigen consists of chains of repeating oligosaccharide subunits, usually composed of two to seven sugars often with additional chemical modifications (15); currently, 182 O groups and 53 flagellar antigen types have been described by phenotypic identification (14, 16). The genes encoding O-antigen biosynthesis are organized in clusters that are flanked by colanic acid biosynthesis gene cluster (*wca* genes) and histidine (*his*) biosynthesis operon (15). O-antigen biosynthesis gene clusters typically have a lower GC content (often <40%) than that of the backbone of the *E. coli* chromosome, which is ∼50% GC content (13, 17, 18). These differences suggest that O-antigen biosynthesis gene clusters are exchanged by lateral gene transfer and are under diversifying selection, and therefore a hot spot of recombination (15, 19).

Although its wide distribution in the affected areas, the near disappearance of *E. coli* O104:H4 in Germany after the large outbreak in 2011 was unanticipated. Here, we show that the high virulence ensemble of the O104:H4 outbreak strain remains a threat, but that O-antigen gene exchange has cloaked the pathogen with several new O-antigens.

## Results

### The HUS-associated STEC O181:H4 strain 17-07187 shares a close phylogenetic relationship and similar virulence traits with the O104:H4 outbreak strain

After the large STEC O104:H4 outbreak in 2011, the German National Reference Centre for *Salmonella*and other Bacterial Enteric Pathogens intensified STEC surveillance and analyzed ∼ 8,000 clinical isolates primarily from diarrhea and HUS patients from 2012 to 2020. This strain collection included a stool sample isolate (17–07187) from a 6-year-old girl who had bloody diarrhea and HUS in December 2017. She had not travelled outside her home in Northwest Germany before becoming ill. Serotyping and whole genome sequencing revealed that the strain belonged to an unusual serotype, O181:H4, that had not been previously associated with HUS. The strain had *stx2a* but lacked the LEE pathogenicity island (marker gene *eae*). Furthermore, the strain encoded characteristic EAEC markers including *aatA, aggR,* AAF/I genes and autotransporter protease genes *pic, sigA,* and *sepA* (Fig. 1A, Tab. S1).

**Figure 1:**
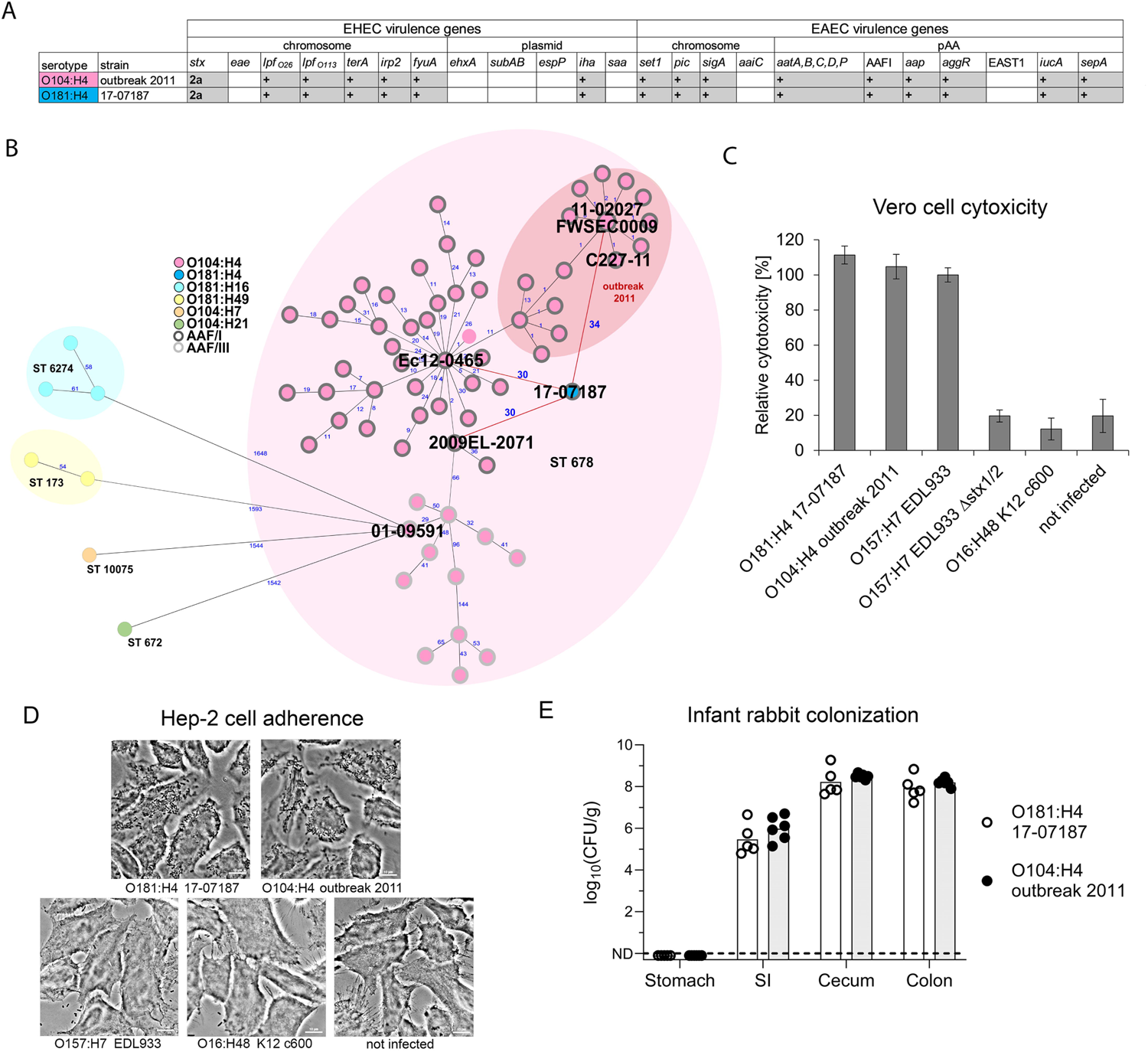
STEC O181:H4 17-07187 shares close phylogenetic relationship and similar virulence traits with the O104:H4 outbreak strain. (A) Overlapping virulence gene profiles of the O104:H4 outbreak strain (FWSEC0009) and the O181:H4 strain 17-07187. (B) Minimal spanning tree of O181:H4 17-07187 and selected O104:H4 strains based on cgMLST involving 2513 alleles confirms their close phylogenetic relationship. Strains representing phylogenetic diversity of O104:H4 isolates from humans with respect to isolation time (from 1998 to 2022) and location were selected. Only one cluster representative was used, except of the 2011 outbreak. O181:H4 17-07187 and O104:H4 strains 2009EL-2071, Ec12-0465, and outbreak strains from 2011 (including strains FWSEC0009, C227-11, and 11-02027) differ by only 30-34 alleles. An allele represents a variant form of a gene. Numbers in blue indicate allelic distances. Serotypes and AAF type are indicated in the key. For complete strain designations see Fig. S1A. (C) Cytotoxicity of O181:H4 17-07187 towards Vero cells and (D) adherence pattern to HEp-2 cells are comparable to the O104:H4 outbreak strain 11-02027. Results are representative of at least two additional experiments. EHEC O157:H7 EDL933 producing Stx1 and Stx2 served as reference in the cytotoxicity assay and was set to 100%. The cytotoxicity results represent the means and standard deviations of triplicate samples (n=3) and are representative of at least two additional experiments. E) Intestinal colonization in infant rabbits inoculated with O104:H4 outbreak strain C227-11 (n=6) or O181:H4 strain 17-07187 (n=5). CFU/g (colony forming units per gram of tissue) denotes concentration of bacteria recovered three days post inoculation from homogenated intestinal tissues. Data points represent individual rabbits from two independent litters split between the two strains. Bars show the geometric mean, SI is small intestine, ND is not detected.

MLST demonstrated that the 17-07187 isolate belonged to the same sequence type (ST678) as the O104:H4 outbreak strain (Table S2) (5, 7, 9). The EnteroBase core genome MLST (cgMLST) scheme (2,513 genes) (20) confirmed the close genomic relationship of this isolate with a panel of O104:H4 strains (Fig. 1B, Fig. S1A). There were only 30-34 allelic distances between this O181:H4 isolate and the O104:H4 outbreak strain (e.g. strains FWSEC0009, C227-11, or 11-02027), and O104:H4 clinical isolates from the Republic of Georgia in 2009 (2009EL-2071) and France in 2012 (Ec12-0465). In contrast, there was considerable allelic distance (AD) to STEC serotypes O181:H16 (ST6274), O181:H49 (ST173), O104:H21 (ST672), and O104:H7 (ST10075) isolated between 2012 and 2019 (AD >1500) (Fig. 1B). Comparison of the virulence gene repertoire of the O181:H4 and the O104:H4 outbreak strain also strongly suggested that they rely on very similar virulence mechanisms (Fig. 1A). Indeed, both strains had comparable Stx-related cytotoxicity (Fig. 1C), exhibited a characteristic enteroaggregative adherence pattern (Fig. 1D), and colonized intestinal tissues, particularly the cecum and colon, similarly during *in vivo* infant rabbit infections (Fig. 1E). Thus, STEC O181:H4 and O104:H4 isolates share marked genomic similarity and virulence-associated genomic and phenotypic traits.

### The genomes of the STEC O181:H4 strain 17-07187 and the O104:H4 outbreak strain mainly differ in their O-antigen gene clusters and mobile genetic elements

The chromosomes (without plasmids) of the O181:H4 isolate 17-07187 and the O104:H4 outbreak strain FWSEC0009 were very similar (∼99.7% nucleotide identity in the core genome that is shared between these strains). The most striking difference between them were their respective O-antigen gene clusters (OAGC) (Fig. 2A, 2B). Although these two clusters were both situated in the same location in the chromosome, between *galF* and *hisI* (Fig. 2B), their gene contents and organization were very different. Furthermore, their respective GC contents, 36.8% for O181 and 37% for O104, differed from the chromosome GC content (∼50.7%), highlighting the likely role of lateral gene transfer in driving OAGC exchange. Fourteen potential prophage regions are present in the O181:H4 isolate and 16 in the O104:H4 outbreak strain (Fig. 2A, Table S3). Eleven of the fourteen prophages exhibited substantial sequence identity (83-99.9%) with their O104:H4 counterparts and importantly, the *stx2-*encoding prophages were nearly identical (∼99.9% nucleotide identity) (Fig. 2C, Table S4). Both *stx* phages are inserted into the tryptophan repressor binding protein gene *wrbA*.

**Figure 2:**
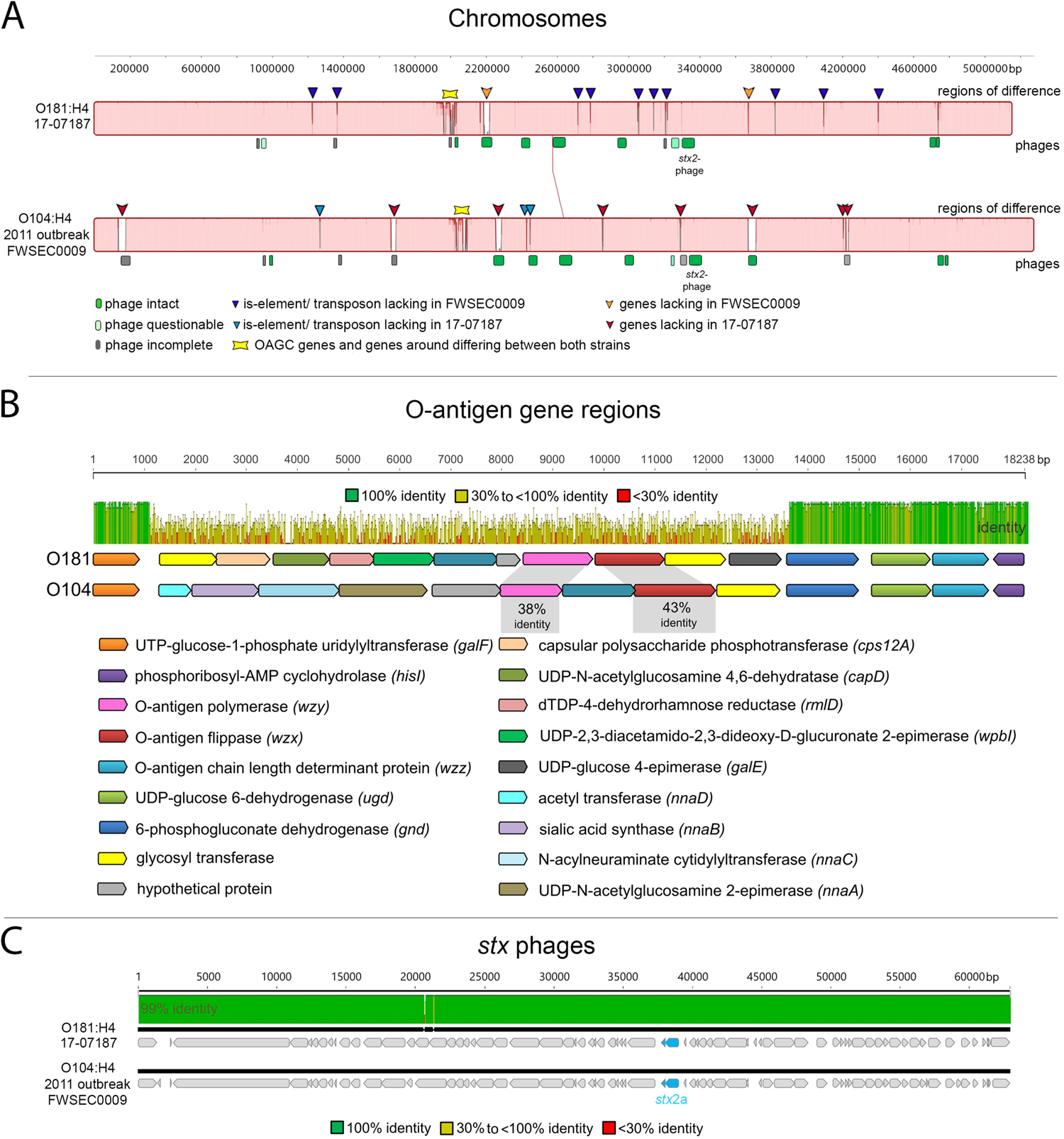
O181:H4 17-07187 and O104:H4 outbreak strain genomes mainly differ in OAGCs and mobile genetic elements. (A) Whole chromosome MAUVE alignment of O181:H4 17-07187 and the O104:H4 outbreak strain FWSEC0009 highlighting mobile genetic elements and differences in phage regions, IS-/ transposon elements, and OAGC regions. (B) OAGCs of O181:H4 17-07187 and the O104:H4 FWSEC0009 outbreak strain are flanked by homologous upstream (*galF*) and downstream (*gnd* to *hisI*) regions. MAFFT alignment shows that the regions in between *galF* and *gnd* are very different. (C) MAFFT alignment of the *stx2a*-encoding phages in O181:H4 17-07187 and the outbreak strain O104:H4 outbreak strain FWSEC0009 shows that they are very similar (99,9% nucleotide identity).

The genome of the O181:H4 isolate included three plasmids of ∼81kb, ∼76kb, and ∼63kb (Table S2). The largest O181:H4 plasmid (pEc17-07187-1) was an incompatibility group I1 (IncI1) plasmid that showed only partial homology to the ESBL resistance plasmid of the O104:H4 outbreak strain (∼75% nucleotide identity) but was very similar (98% nucleotide identity) to pHUSEC41-1 of STEC O104:H4 HUSEC41 from 2001 (92kb) (Fig. S1B) (3, 21). Both of these plasmids encode the *pilI-V* genes for thin pili. Unlike pHUSEC041-1, the O181:H4 pEc17-07187-1 did not contain antibiotic resistance genes (Fig. S1B, Tab. S5). O181:H4 plasmid (pEc17-07187-2/pAA) (76 kb) was very similar (99.8% nucleotide identity) to the O104:H4 outbreak strain pAA-EA11 and harbored virulence associated loci including *aggA/B/C/D*, which encode the AAF/I (Fig. 1A, Fig. S1C, Tab. S6) (7, 11). O181:H4 plasmid pEc17-07187-3 (63 kb) was not found in the O104:H4 outbreak strain; instead, it showed similarity to DHA plasmids of several enterobacteria coding for AmpC β-lactamase (22). However, unlike the DHA plasmids, resistance determinants were not present in the O181:H4 isolate (Fig. S1D, Tab. S7). Together these observations reveal the striking similarity of the chromosomes of the O181:H4 isolate and the O104:H4 outbreak strains and that their chief differences are confined to hot spots of recombination, i.e. their O-antigen gene clusters, and to mobile genetic elements, particularly their plasmids.

### Additional recent global isolates of serotype O181:H4 and five other O groups belong to ST678

Next, we identified 158 genomes belonging to ST678 in EnteroBase (20), which contains ∼202,200 *E. coli* genomes (as of April 11th, 2022). For a subset, serotype identification was not available, and for these cases, serotype was predicted based on the available genomic information (16). 123 of the ST678 strains were O104:H4, however, 18 additional O181:H4 genomes of ST678 were found. Furthermore, seven O127:H4, three O131:H4, and one each of O69:H4 and OX13:H4 were identified (Fig. 3A, Tab. S1). We categorized these as non-O104:H4 ST678 strains. Additionally, based on a close phylogenetic relationship and single difference in MLST alleles, we added three non-ST678 strains to the non-O104:H4 ST678 category: two O181:H4 strains (1472912 and 1472968) and one strain of the new and provisionally assigned genoserotype OgN-RKI9:H4 (strain 608450) (Fig. 3A, Tab. S1).

**Figure 3:**
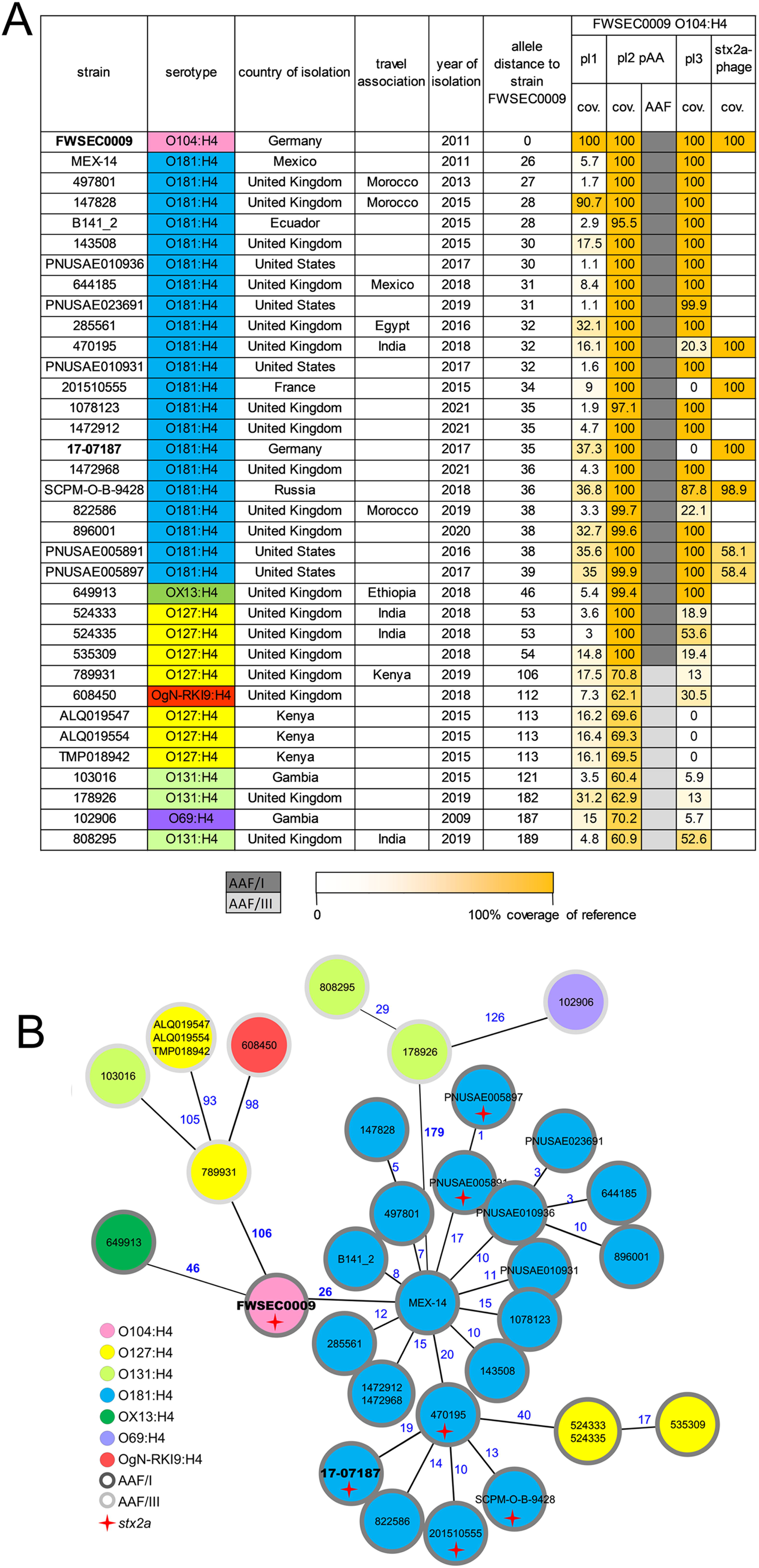
Recent global isolates of serotype O181:H4 and five other O groups belong to ST678. (A) Summary of serotypes, isolation dates, along with relatedness of genomes (cgMLST-based allelic distances) and mobile genetic elements (plasmids and *stx* phage) of the 34 non-O104:H4 ST678 (ST12598, ST12610) clinical strains (found on EnteroBase) compared to the O104:H4 outbreak strain FSWEC0009. (B) Serotypes of the non-O104:H4 ST678 strains and the O104:H4 outbreak strain FSWEC0009 in a minimal spanning tree based on cgMLST. Numbers in blue indicate allelic distances. Although, phylogenetically very close to ST678 strains, O181:H4 strains 1472912 and 1472968 in fact belong to ST12610 due to a point mutation in *icd* and OgN-RKI9:H4 strain 608450 to ST12598 due to a point mutation in *recA*.

cgMLST confirmed the close genetic relationship (AD ∼50) between AAF/I gene positive O104:H4 strains including the outbreak strain (FWSEC0009), and the 21 O181:H4 strains, the OX13:H4 strain, and three of the seven O127:H4 strains (Fig. 3A, 3B). All of these strains showed AAF/I which were also found in the 2011 outbreak strain (5, 7). Nine strains (four O127:H4, three O131:H4, one each of O69:H4 and OgN-RKI9:H4) had higher allelic distances up to 189. In contrast, these nine strains encoded AAF/III genes (Fig. 3A, 3B). The 25 AAF/I positive non-O104:H4 ST678 strains were all isolated in or after 2011 and the majority was associated with diarrheal disease and a subset of six O181:H4 strains harbored *stx2a* (Tab. S1, Fig. 3A, 3B, Fig. S2A). Interestingly, four of these six genomes (including 17-07187 from Germany) shared a very similar *stx* phage with the O104:H4 outbreak strain, but two of the six prophages were more distinct (Fig. 3A). AAF/I-positive strains shared regions with a higher relatedness than those positive for AAF/III with the pAA plasmid of the 2011 outbreak strain (Fig. 3A).

The virulence gene profiles of the 34 non-O104 ST678 strains were generally similar to the O104:H4 outbreak strain; however, there were a few differences. Specifically, *sepA* was exclusively found in AAF/I-positive strains and EAST1 was present in all AAF/III-positive strains and in only three of the AAF/I-positive isolates (Tab. S8).

The 34 non-O104:H4 ST678 strains were isolated in countries of Europe, Africa, and North and South America. In addition, several were from individuals with a travel history that might link these to East Asia (Fig. 3A, S2B and C). Together, these observations show that ST678 *E. coli* strains are found among seven different *E. coli* serotypes that have been linked to diarrheal disease on several continents.

### Phylogenomic analyses of ST678 *E. coli* suggest occurrence of multiple O-antigen gene exchange events

The O-antigen gene clusters encoding the six O groups in the 34 non-O104:H4 ST678 strains are found in the same chromosomal location as the 2011 O104:H4 outbreak strain but are composed of largely disparate genes (Fig. 4A). These clusters also have a distinct GC content (36.8-42.1%) from the backbone genome (50.5-50.7%), suggesting that they were acquired by horizontal gene exchange. To explore the phylogenetic relationships among the 34 non-O104:H4 ST678 strains and a set of O104:H4 strains, their shared single nucleotide polymorphisms (SNPs) were analyzed using the O104:H4 outbreak strain FWSEC0009 as a reference. A positive correlation (R=0.81; R^2^=0,66) was found between isolation time and genetic divergence (Fig. S2D) which supports that mutations have accumulated in a clock-like fashion without notable outliers. A phylogenetic analysis shows that AAF/I positive strains within ST678 are closely related to each other, clustering in clade I, whereas AAF/III positive strains are more diverse and form more basal branches in a rooted maximum-likelihood phylogenetic tree (Fig. 4B). This structure of the phylogeny suggests that clade I strains were derived from an AAF/III-positive precursor. Within clade I, subclades Ia and Ib contain some non-O104:H4 strains. Clade Ia is composed of O104:H4 non-outbreak strains isolated from 2015-2021 in the United Kingdom and in Kenya and the 2018 OX13:H4 strain that was associated with travel to Ethiopia. Clade Ib contains all 21 O181:H4 strains and three O127:H4 strains isolated from 2011 to 2021 from diverse continents. Since the majority of ST678 strains are serotype O104:H4, including basal phylogenetic branches and most isolates in clade I (Fig. 4B), it is parsimonious to assume that subclade Ib emerged from a O104:H4 predecessor, by replacing O104 antigen genes by O181 antigen genes. Similarly, our phylogenetic analysis suggests that O127:H4 strains within clade Ib arose from an O181:H4 precursor. Together these observations indicate that O-antigen gene cluster exchange has occurred repeatedly within ST678 strains.

**Figure 4:**
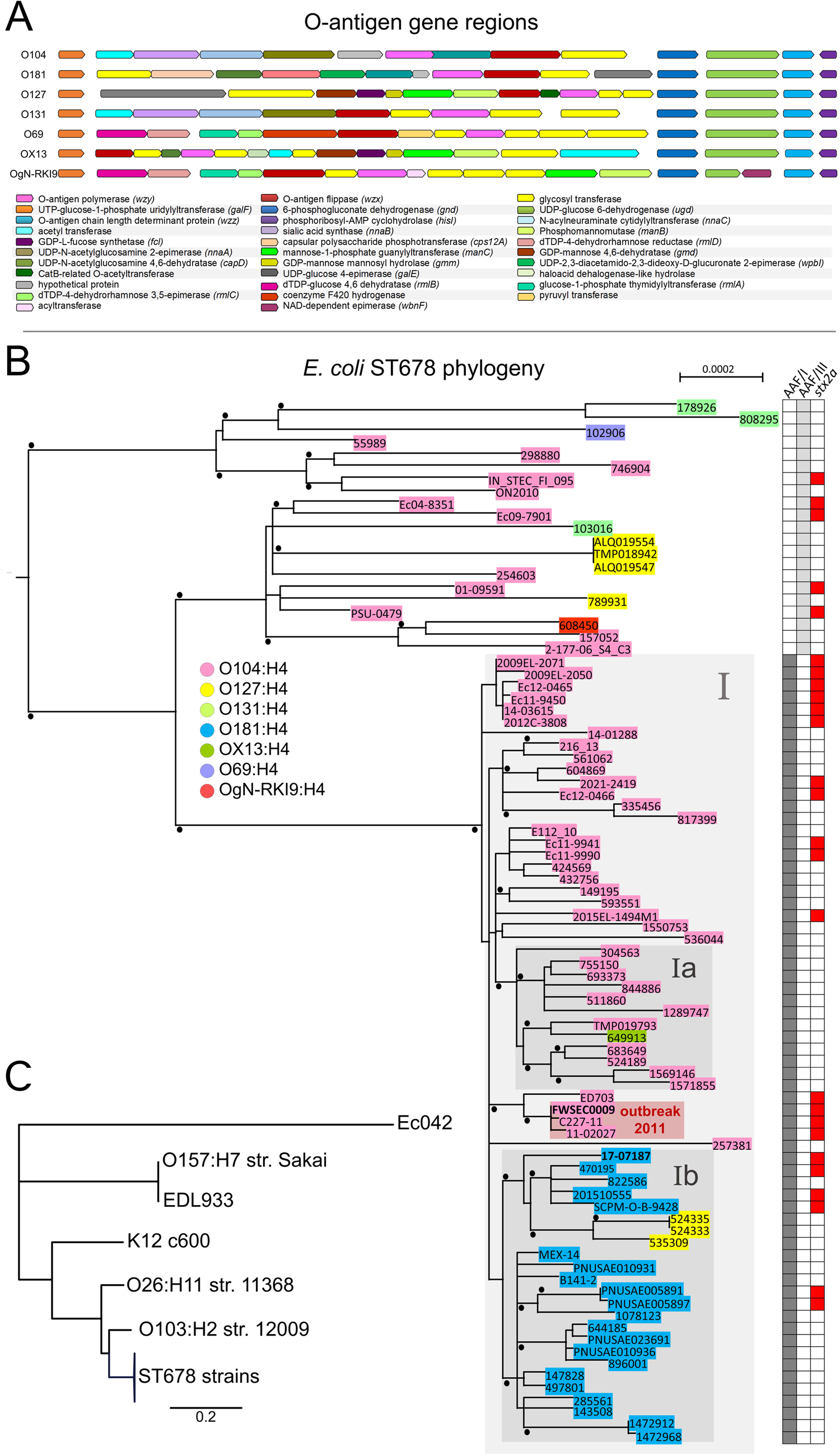
O-antigen gene clusters and phylogeny within *E. coli* ST678. (A) O-antigen gene clusters in ST678 strains differ in their gene contents and organization, yet they are flanked by homologous regions starting upstream by *galF* and downstream by *gnd* to *hisI*, respectively. (B) Maximum-likelihood phylogenetic tree generated by PhyML based on a recombination-corrected alignment of genome-wide polymorphic sites. Strains selected for representation in Fig. 1B and Fig. 3 were combined and O104:H4 2011 outbreak strain FWSEC0009 served as reference for read mapping. The tree was rooted with an outgroup consisting of *E. coli* strains K12 c600, Ec042, EDL933, O157:H7 strain Sakai, O26:H11 strain 11368, and O103:H2 strain 12009. Genome sequences from outgroup *E. coli* isolates had been included in the phylogenetic analysis and for clarity are not shown in Fig. 4B (large phylogenetic distance and associated limited resolution of the clades of interest) but in Fig. 4C. Bootstrap values >90% are indicated as black dots. Serotypes are as indicated in the key. Heatmap on the right indicates the presence (filled box) or absence (white box) of genes for AFF/I, AFF/III, and *stx2a*. Scale bar refers to a phylogenetic distance of 0.0002 nucleotide substitutions per site. Grey boxes indicate clades. The box in light red highlights 2011 outbreak strains. (C) Maximum-likelihood phylogenetic tree with additional outgroup *E. coli* isolates and strains represented in Fig. 4B as collapsed tree branch designated as ST678 strains.

## Discussion

The *E. coli* O104:H4 outbreak in Germany in the early summer of 2011 was a public health emergency; however, this serotype was rarely isolated as a cause of HUS after the epidemic subsided. Nevertheless, our findings suggest that the unusual set of virulence factors that characterized this Shiga toxin 2-producing EAEC strain remains a threat to human health. Serotype conversion has cloaked this highly virulent genotype with several O-antigens, including O181, O127, and OX13, which are present both in *stx* positive and *stx* negative disease-linked isolates closely related to the O104:H4 outbreak strain. We found non-O104:H4 ST678 strains from a variety of countries in Europe, Africa, and the Americas and several of the cases were associated with travel to Africa and Asia, suggesting these virulence-associated strains are globally distributed. Like the O104:H4 outbreak strain, the AAF/I positive ST678 strains had similar chromosomes, similar pAA-linked virulence genes, as well as virulence factors including the SPATEs (5, 7, 8, 12). The most salient difference of the chromosomes of these strains with that of the outbreak strain were their respective O-antigen gene clusters and other mobile genetic elements. Thus, horizontal exchange of these clusters appears to have been a critical step in the evolution of these new pathogenic serotypes, some of which were linked to HUS or bloody diarrhea.

The prime example uncovered here is the likely derivation of O181:H4 pathogens from an O104:H4 precursor via O-antigen gene exchange. Among the 21 O181:H4 strains, six harbored a *stx2a*-encoding prophage, including HUS-linked strain 17-07187. In four of these strains, the *stx2a* prophage was very similar to the *stx2a* prophage of the 2011 outbreak strain (Fig 3A), suggesting that the O-antigen exchange was a more recent event in their evolution compared to the acquisition of the Shiga toxin-encoding prophage. The absence of *stx* prophages in 15 of the O181:H4 isolates may be due to a lack of *stx* prophage acquisition or have resulted from loss of their *stx* prophages, which is well documented in STEC in other serotypes (23). Thus, in addition to O-antigen exchange, on-going horizontal transmission of mobile genetic elements, such as *stx* phages, has contributed to diversification of diarrheagenic ST678.

O-antigen gene clusters of Gram-negative bacteria are hot spots for diversifying selection and recombination events (15, 19). Serotype conversion by lateral exchange of OAGCs has played an important role in the evolution of enteric pathogens. For example, *Vibrio cholerae* serogroup O139, which arose by exchange of O1 and O139 OAGCs, transiently replaced the dominant O1 group as the cause of cholera in 1992-93 (13, 24). Our discovery of seven distinct O-groups that share the flagellar H4 antigen and a highly similar virulence-linked genetic makeup provides a compelling example of the role of O-antigen exchange in the diversification of diarrheagenic *E. coli*. In addition, STEC O157:H7 is thought to have arisen from an enteropathogenic *E. coli* (EPEC)-like O55:H7 strain which initially acquired *stx* via phage transduction and subsequently acquired the O157 O group by exchange of the O55 with the O157 OAGC (26). Also, an O182-O156 switch is thought to have occurred in STEC strains persistently infecting cattle (18). Consequently, different OAGs may be found in highly related genotypes (27, 28).

We can only speculate about the conditions driving OAGC exchange among ST678 strains. Shiga toxin-producing O104:H4 EAEC strains, such as the 2011 outbreak strain, are considered human-restricted pathogens and have not been isolated from animals, such as cattle (29). However, interestingly a recent description highlighted the detection of an STEC O104:H4 strain in pork (30). It is possible that the OAGC exchange occurred in a human host, where the human intestinal microbiome may contain *E. coli* strains of O groups, such as O181 and O127, that could have donated their OAGC to an O104:H4 recipient by some means of horizontal exchange. Also, epidemiological investigations suggest that fenugreek sprouts were the food source that initiated the STEC O104:H4 epidemic; therefore plants colonized by microbiota may be another possible site for OAGC exchange (4).

## Conclusion

In conclusion, our study highlights how lateral exchange of O-antigen gene clusters can lead to the rapid diversification of a globally important pathogen. Further, an important clinical implication of these findings is that serotype surveillance cannot be used as a simple proxy for strain virulence and needs to be complemented by virulence gene or genome analysis. Surveillance to uncover how highly virulent strains may reemerge and spread in new O-antigen outfits is warranted.

## Material and Methods

### Study strains

In the context of intensified STEC surveillance in Germany, clinical isolates were collected at the National Reference Center for *Salmonella* and other Bacterial Enteric Pathogens and analyzed for serotype, *stx* and subtypes, *eaeA*, *hlyA* and *aatA* as described (31). Strains were grown on nutrient agar (Oxoid GmbH, Germany), Luria Bertani (LB) broth, or enterohemolysin agar (Sifin GmbH, Germany). Genome sequencing was carried out on a subset of strains and further genome sequence data of strains was gathered from EnteroBase and NCBI (Table S1).

### Whole genome sequencing (WGS)

Long read whole genome sequencing of O181:H4 strain 17-07187 was performed by GATC Biotech (Konstanz, Germany) using a PacBio RS II sequencer (Pacific Biosciences, USA) and short read genome sequencing of strains 12-04810, 14-03615, 14-01288, 16-00596, 16-01499, 16-05332, 17-01774, 17-00416, 17-07187 and 19-02696 (Table S1) on an Illumina MiSeq and HiSeq 1500 benchtop sequencer.

### Bioinformatics analyses

*De novo* assembly of the PacBio sequence data (103-fold average coverage) was performed by GATC utilizing HGAP3 (Pacific Biosciences, USA). Polishing of the assembled genome and plasmids was performed with Illumina short reads by Pilon (version 1.22) (32). Quality control and trimming of MiSeq raw reads with subsequent detection of serotype and virulence genes was performed as described (16). Genomic comparisons were carried out using MAUVE (version: 1.1.1) and MAFFT (version 1.3.7) as plugin in Geneious (version: 11.1.5; Biomatters Ltd) (33, 34). Gaps within the MAFFT-alignment were excluded using the Geneious “mask alignment” function. Ridom SeqSphere+ (version:7.2.0, Ridom GmbH, Germany) was used to determine MLST Warwick sequence types and to create minimal spanning trees based on 2513 allele targets (here genes) from the *E. coli* cgMLST EnteroBase scheme with pairwise ignoring missing values from genome assemblies (20). Different variants of the cgMLST genes among different genomes are represented as allelic distances (AD). Phage prediction was carried out by analysing the genome sequences with PHASTER (35). RAST was used for CDS annotation (36).

### SNP-based alignment and maximum-likelihood based phylogenetic tree

Mapping of sequencing reads, generation of consensus sequences, and alignment calculation was performed using the BatchMap pipeline with FWSEC0009 genome sequence as reference (37). Gubbins (version 3.2.1) was used to identify loci containing elevated densities of base substitutions (putative recombinations) within the alignment while concurrently constructing an alignment and phylogeny (RAxML-tree) based on the putative point mutations (SNPs) outside of these regions (38). Alignment generated by Gubbins was used to create a maximum-likelihood based phylogenetic tree by PhyML 3.3.20180214 (Geneious plugin, substitution model: HKY85, 100 bootstraps) (39).

### Temporal signal and ‘clocklikeness’ of molecular phylogenies

TempEst was used to analyze the RAxML tree generated by Gubbins in conjunction with collection year data to validate the molecular-clock assumption (40). Best-fitting root was chosen for linear regression analyses.

### Cytotoxicity, adherence, and infection assays

Viability of Vero cells after inoculation with diluted bacterial culture supernatants (1:200) was examined using 3-[4,5-dimethylthiazole-2-yl]-2,5-diphenyltetrazolium bromide (MTT) (5). Bacteria and Vero cells were prepared as described (31). Adherence to HEp-2 cells was performed as described (37). For infant rabbit infection assays, litters of mixed gender 2-day-old New Zealand White infant rabbits with the lactating doe were acquired from Charles River (Canada, strain code 052). Infant rabbits were orogastrically inoculated on the day of arrival with 10^9^ CFU of Streptomycin-resistant strains O104:H4 C227-11 and O181:H4 17-07187 suspended in 500µl 2.5% sodium bicarbonate (pH9) using a size 4 French catheter as described previously, except that no ranitidine was administered (12). No antibiotics were used prior or during infection. Infant rabbits were monitored for signs of illness and euthanized three days (68-72 hours) post infection. Tissue samples taken from the stomach, small intestine, cecum, and colon were homogenized in sterile PBS using a minibeadbeater-16 (Biospec Products, Inc.) and CFU determined by serial dilution, and plating on LB media containing 200 µg/ml streptomycin (12).

### Ethics statement

Rabbit studies were conducted according to protocols reviewed and approved by Brigham and Women’s Hospital Committee on Animals (IACUC protocol 2016N000334) and Animal Welfare Assurance of Compliance (number A4752-01) in accordance with recommendations in the Guide for the Care and Use of Laboratory Animals of the National Institutes of Health and the Animal Welfare Act of the U.S. Department of Agriculture.

### Data availability

The generated sequences were uploaded to NCBI project: PRJNA833419 (https://www.ncbi.nlm.nih.gov/bioproject/?term=PRJNA833419).

## Acknowledgements

We thank Ute Strutz, Thomas Garn, Tobias Größl, and Karsten Großhennig for excellent technical assistance (Robert Koch Institute, Wernigerode, Germany). We also acknowledge the MF1 bioinformatics and MF2 genome sequencing unit of the Robert Koch Institute (Berlin) for the support of bioinformatic analysis and Illumina MiSeq and MinION sequencing. We thank members of the Waldor and Flieger groups for helpful comments and discussions during the study. AFl received funding from the German Ministry of Health (IGS-ZOO grant number 2518FSB706) and AFl and AFr from the intensified molecular surveillance initiative of the Robert Koch-Institute. IWC received funding from the National Institute of Diabetes and Digestive and Kidney Diseases of the National Institutes of Health (Award Number 5 T32DK007477-37).

## Author Contributions

conceptualized the study: C.L., A.Fl.

supervised the study: A.Fl.

conceived and designed the experiments: C.L., A.Fr., I.W.C., U.N., Y.H.G., M.K.W., A.Fl.

acquired data: C.L., A.Fr., I.W.C., C.J., P.S., N.S., F.-X.W., A.Fl.

conducted the experiments: C.L., I.W.C, A.Fl.

analyzed the data: C.L., A.Fr., I.W.C., C.J., P.S., N.S., F.-X.W., U.N., Y.H.G., A.Fl.

interpreted the results: C.L., A.Fr., I.W.C., C.J., P.S., N.S., F.-X.W., U.N., Y.H.G., M.K.W., A.Fl.

drafted the manuscript: C.L., M.K.W., A.Fl.

revised the manuscript: C.L., A.Fr., I.W.C., C.J., P.S., N.S., F.-X.W., U.N., Y.H.G., M.K.W., A.Fl.

approved the final version: C.L., A.Fr., I.W.C., C.J., P.S., N.S., F.-X.W., U.N., Y.H.G., M.K.W., A.Fl.

