## Supplementary material for "O-antigen diversification masks identification of highly pathogenic STEC O104:H4-like strains": Supplemebtal Table S2

**Table S2:** Genome characteristics of HUS-associated STEC O181:H4 17-07187 and the 2011 STEC O104:H4 outbreak strain FWSEC0009. CDS = coding sequences: determined by means of RAST annotation (http://rast.nmpdr.org/rast.cgi)

| Strain | 17-07187 | FWSEC0009 |
| --- | --- | --- |
| Serotype | **O181:H4** | **O104:H4** |
| MLST | ST678 | ST678 |
| Epidemiology | Sporadic case | 2011 EHEC outbreak |
| Year/Country of isolation | 2017 / Germany | 2011 / Germany |
| Clinics | HUS | Not stated |
| Accession | NCBI: project: PRJNA833419 | NCBI: CP031902 |
| chromosome size | 5 155 820 bp | 5 277 234 bp |
| GC content% | 50.7 | 50.7 |
| chromosome CDS^#^ | 5137 | 5270 |
| plasmid 1 | 80 851 bp | 88 545 bp |
| plasmid 1 CDS | 117 | 114 |
| plasmid 2/pAA | 75 597 bp | 74 217 bp |
| plasmid 2/pAA CDS | 128 | 126 |
| plasmid 3 | 63 387 bp | 1 549 bp |
| plasmid 3 CDS | 96 | 3 |
