## Supplementary figures and images for "O-antigen diversification masks identification of highly pathogenic STEC O104:H4-like strains"

### Supplemebtal Figure S1

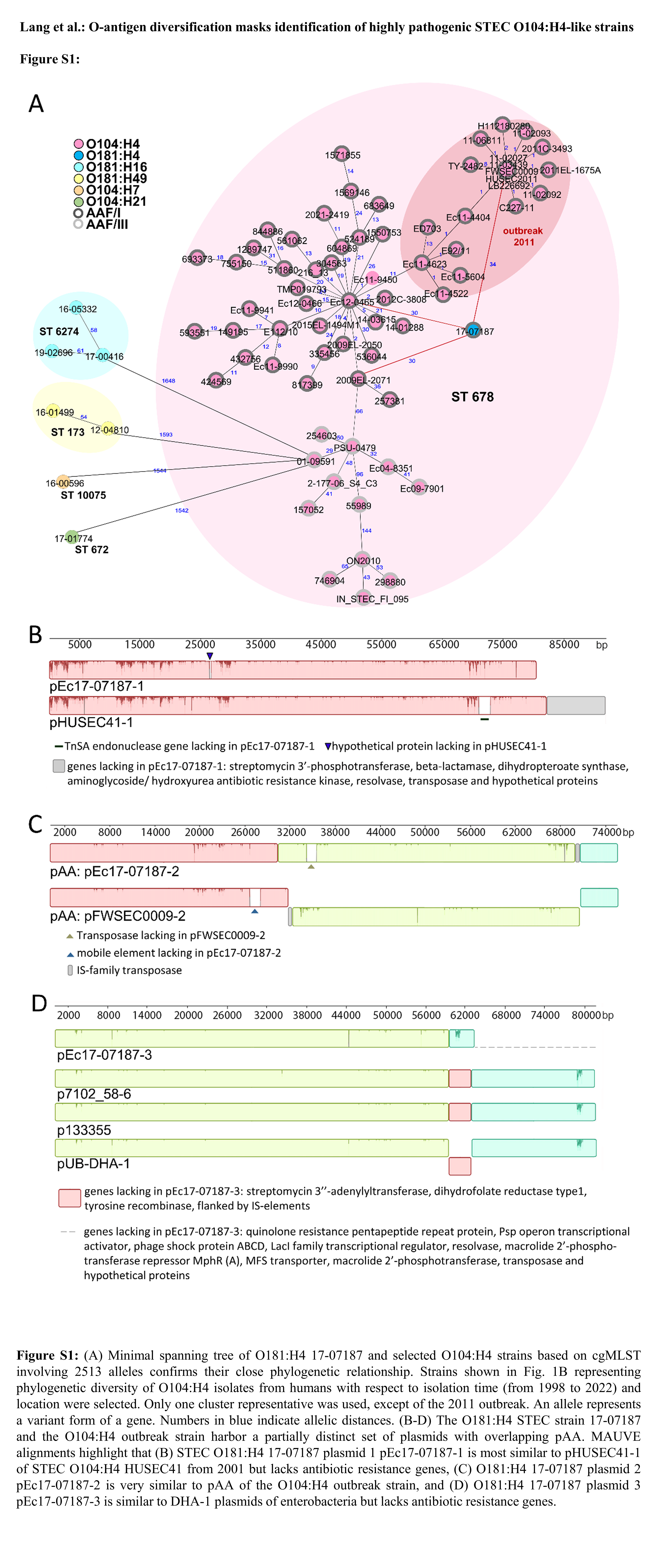

### Supplemebtal Figure S2

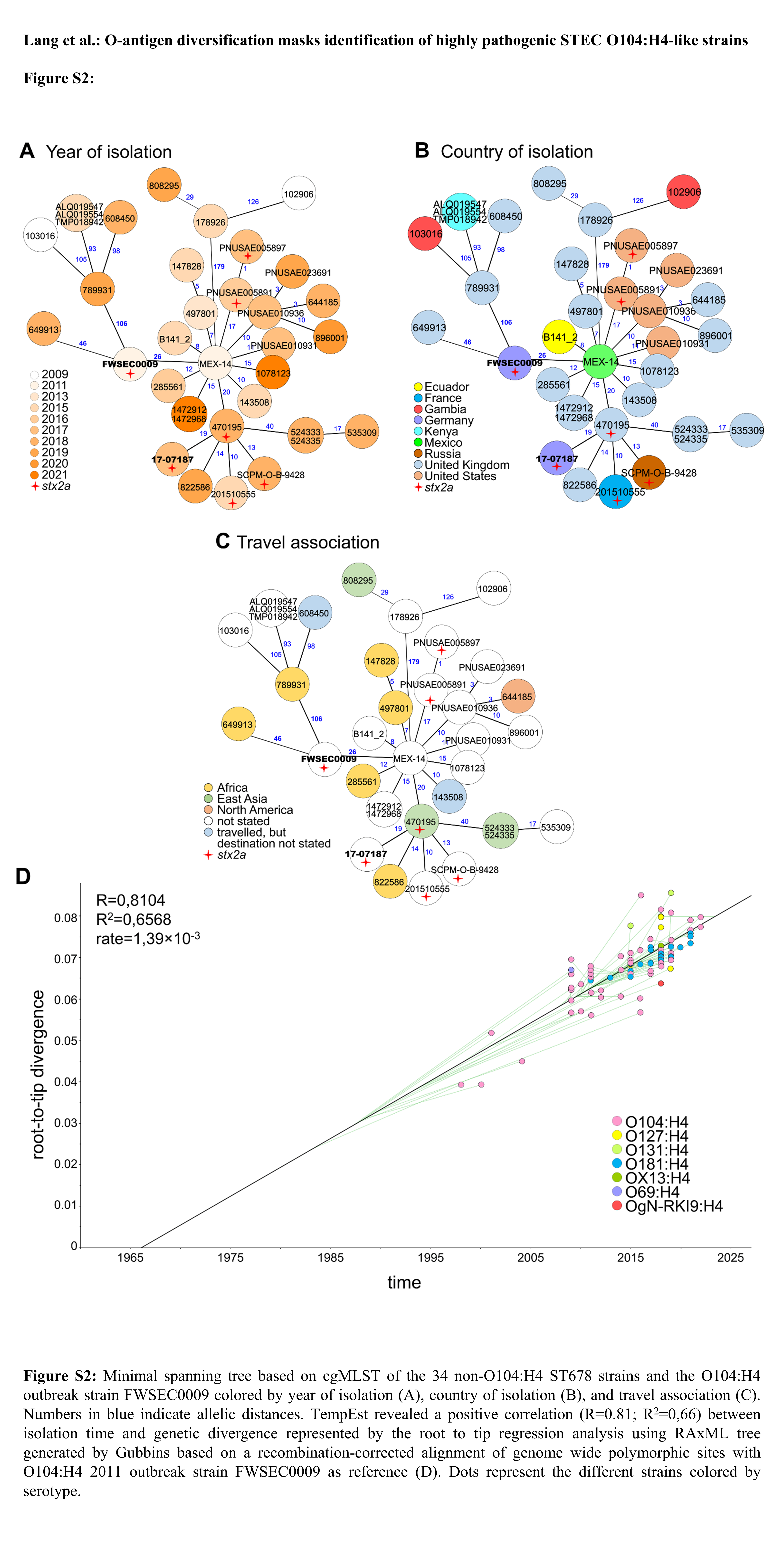
